# PERSONALIZED BIOMARKERS OF MULTISCALE FUNCTIONAL ALTERATIONS IN TEMPORAL LOBE EPILEPSY

**DOI:** 10.1101/2025.04.24.650457

**Authors:** Ke Xie, Ella Sahlas, Alexander Ngo, Judy Chen, Thaera Arafat, Jessica Royer, Yigu Zhou, Raúl Rodríguez-Cruces, Arielle Dascal, Benoit Caldairou, Fatemeh Fadaie, Alexander Barnett, Samantha Audrain, Sara Larivière, Lorenzo Caciagli, Raluca Pana, Alexander G. Weil, Christophe Grova, Birgit Frauscher, Dewi V. Schrader, Zhiqiang Zhang, Luis Concha, Andrea Bernasconi, Neda Bernasconi, Boris C. Bernhardt

## Abstract

Temporal lobe epilepsy (TLE) presents with substantial inter-patient variability in clinical and neuroimaging manifestations. This multicenter study examined inter-individual differences in spatial patterns of intrinsic brain function in TLE using normative modeling at multiple spatial scales and evaluated the effectiveness of individual functional deviations for clinical diagnosis and postsurgical outcome prediction. We analyzed multimodal MRI data on 298 healthy controls, 282 TLE patients, and 45 disease controls with extratemporal epilepsy. Cortical function was profiled at local, regional, and global scales using brain signal variability, regional homogeneity, and node strength. We estimated patient-specific *W*-score maps to index deviations from normative metrics. Compared to healthy controls, patients with TLE showed considerable variations in patterns of functional alterations across the cortex, with the highest overlap in the ipsilateral mesiotemporal regions. Connectome-based simulation revealed the paralimbic and medial default mode regions as key disease epicenters. Functional changes were primarily underpinned by superficial white matter anomalies. Supervised pattern learning achieved classification AUCs of 0.76 for TLE versus disease controls, 0.74 for left versus right TLE, and 0.63 for seizure-free versus non-seizure-free TLE, with greater contralateral temporal functional deviations correlating with unfavorable postsurgical seizure outcome. Our findings reveal the heterogeneous impact of TLE on intrinsic cortical function. These biomarkers hold promise for clinical translation, guiding precision therapeutics and enhancing presurgical decision-making in TLE.

## 1. Introduction

Temporal lobe epilepsy (TLE) is the most prevalent pharmaco-resistant focal epilepsy in adults. Magnetic resonance imaging (MRI) serves as a powerful tool for *in vivo* profiling of brain structure and function, and has been widely applied in the diagnosis and management of TLE.^1-5^ An growing number of studies highlight the substantial variability in the distribution and severity of cortex-wide structural and functional anomalies among individuals with TLE,^6-10^ underscoring the complexity of this disorder. Addressing heterogeneity has been recognized as an essential element in understanding neurobiological mechanisms and in developing robust quantitative MRI biomarkers to support precision diagnosis and prognosis in TLE.^6,7^

Quantitative MRI analyses of TLE have relied on classic case-control designs, which compare patient groups to healthy controls to identify group-level common patterns of brain abnormalities. While this method has helped identify hallmarks of TLE, such as mesiotemporal atrophy and hyperexcitability,^3,4,11^ it provides limited insights into the variability of biological mechanisms across individuals, as it does not pinpoint the precise anatomical loci of pathological changes in any single patient. New paradigms are needed to better model and explain this individual variation. Recent studies based on clustering techniques have revealed subtypes of TLE with distinct patterns of grey and white matter abnormalities associated with different cognitive performance and clinical characteristics,^6,12-14^ reinforcing the need to account for disease heterogeneity. Despite growing research on functional disruptions in TLE, however, most analyses remain at a group-level, hindering the progress toward patientcentered precision medicine.

In this study, we utilized a multicenter, multimodal MRI dataset to profile intrinsic functional imbalances in TLE across multiple spatial scales at the single-patient level using normative modeling. MRI-based normative modeling is an emerging family of techniques that shifts the focus from group average to intra-cohort variability, enabling the identification of patient-specific alteration.^15-19^ This technique models the relationship between MRI features and biological variables (*e*.*g*., age, sex) to establish normative percentiles of variation for each brain region in healthy populations.^20-22^ Each patient is then mapped onto the normative percentile charts to determine deviations (*i*.*e*., *W*-score), capturing where and how an individual differs from the healthy range. Unlike conventional *z*-normalization that relies solely on the average and standard deviation of healthy controls,^23^ normative modeling provides age-and sex-specific inferences, accounting for potential aging effects and sex differences. This technique, widely applied in neurological and psychiatric disorders,^24-29^ has shown promise in detecting individual patterns of variations in brain structure and function, motivating its application to TLE.

Here, we sought to characterize brain functional deviations in individuals with TLE at multiple scales of MRI analysis, to determine the extent to which spatially heterogeneous deviations may accumulate within similar brain regions or distributed brain networks. We conducted one of the largest restingstate functional (rs-fMRI) studies in TLE to date, including 298 healthy controls, 282 TLE patients, and 45 disease controls. Specifically, we first estimated region-specific *W*-scores to quantify functional deviations from normative models at the individual patient level and computed the prevalence of extreme deviations at the group level. Next, we investigated how global brain connectivity shapes local dysfunction and identified potential disease epicenters using a structural connectome-based model that simulates the spread of pathology across networks. As TLE patients are often associated with widespread grey and white matter disruptions, we then examined the relationship between the anatomical distribution of functional and structural deviations. Finally, we evaluated the effectiveness of patient-specific functional deviation scores for diagnosing TLE (versus disease controls), identifying seizure onset zone laterality, and predicting postsurgical seizure outcome using supervised machine learning techniques.

## 2. Results

## 2.1 Data Samples

We analyzed 625 participants, including 282 patients with TLE (133 males; mean ± SD age = 32.3 ± 10.0 years, range: 18–64 years]), 298 healthy individuals (152 males; 29.3 ± 8.0 years [18–60 years]) and 45 disease controls with extratemporal focal cortical dysplasia (FCD) (23 males; 27.2 ± 7.9 years [18–54 years]). Participants were selected from 4 independent datasets of 3 epilepsy centers: (i) Montreal Neurological Institute-Hospital (*MICA-MICs*: TLE/healthy controls/FCD = 57/100/17; *NOEL*: 72/42/28),^30^ (ii) Universidad Nacional Autónoma de México (*EpiC*: 29/34);^14^ and (iii) Jinling Hospital (*Nanj*: 124/122).^31^ Details on subject inclusion are provided in the Methods. Detailed demographic and clinical information for each dataset is provided in **Table 1** and **Figure 1A**.

**Table 1.** Demographic and clinical information.

| dataset<br>(n) | group<br>(n) | age | sex<br>(M/F) | focus<br>(L/R) | age at<br>onset | disease<br>duration | HS<br>(%) | DRE<br>(%) | surgery<br>(Engel I) |
| --- | --- | --- | --- | --- | --- | --- | --- | --- | --- |
| *all*<br>(625) | HC<br>(298) | 29.3±8.0<br>(18–60) | 152/146 | – | – | – | – | – | – |
| | TLE<br>(282) | 32.3±10.0<br>(18–64) | 133/149 | 147/135 | 17.8±10.7<br>(0.3–60) | 14.4±11.0<br>(0.8–49) | 192<br>(68.1%) | 214<br>(75.9%) | 99 (74) |
| | FCD<br>(45) | 27.2±7.9<br>(18–54) | 23/22 | 19/26 | 11.5±6.9<br>(0.5–27) | 15.9±10.3<br>(0.5–48) | – | 40<br>(88.9%) | 36 (25) |
| *MICs*<br>(174) | HC<br>(100) | 32.1±7.4<br>(20–60) | 53/47 | – | – | – | – | – | – |
| | TLE<br>(57) | 36.8±11.3<br>(20–64) | 31/26 | 35/22 | 21.1±13.1<br>(0.5–60) | 15.7±11.1<br>(1–45) | 24<br>(42.1%) | 50<br>(87.7%) | 15 (12) |
| | FCD<br>(17) | 26.9±7.5<br>(19–42) | 7/10 | 6/11 | 11.5±6.0<br>(0.7–22) | 16.0±9.5<br>(0.5–34) | – | 12<br>(70.6%) | 8 (5) |
| *EpiC*<br>(63) | HC<br>(34) | 32.9±11.3<br>(18–57) | 10/24 | – | – | – | – | – | – |
| | TLE<br>(29) | 31.3±11.3<br>(18–58) | 9/20 | 18/11 | 15.0±10.7<br>(1–40) | 16.0±13.1<br>(1–49) | 17<br>(58.6%) | 29<br>(100%) | – |
| *Nanj*<br>(246) | HC<br>(122) | 25.3±5.4<br>(19–40) | 65/57 | – | – | – | – | – | – |
| | TLE<br>(124) | 28.3±7.4<br>(18–48) | 66/58 | 59/65 | 16.8±9.0<br>(0.3–43) | 11.4±8.7<br>(0.8–37) | 113<br>(91.1%) | 63<br>(50.8%) | 35 (26) |
| *NOEL*<br>(142) | HC<br>(42) | 31.5±8.0<br>(20–53) | 24/18 | – | – | – | – | – | – |
| | TLE<br>(72) | 36.0±9.7<br>(19–61) | 27/45 | 35/37 | 17.7±11.2<br>(0.6–51) | 18.0±12.2<br>(1–45) | 38<br>(52.8%) | 72<br>(100%) | 49 (36) |
| | FCD<br>(28) | 27.4±8.3<br>(18–54) | 16/12 | 13/15 | 11.6±7.5<br>(0.5–27) | 15.8±10.9<br>(3–48) | – | 28<br>(100%) | 28 (20) |

Age, age at seizure onset, and disease duration are presented as mean ± SD years. M = male; F = female; L = left; R = right; HS = hippocampal sclerosis; DRE = drug-resistant epilepsy; HC = healthy controls; TLE = temporal lobe epilepsy; FCD = focal cortical dysplasia.

**Figure 1:**
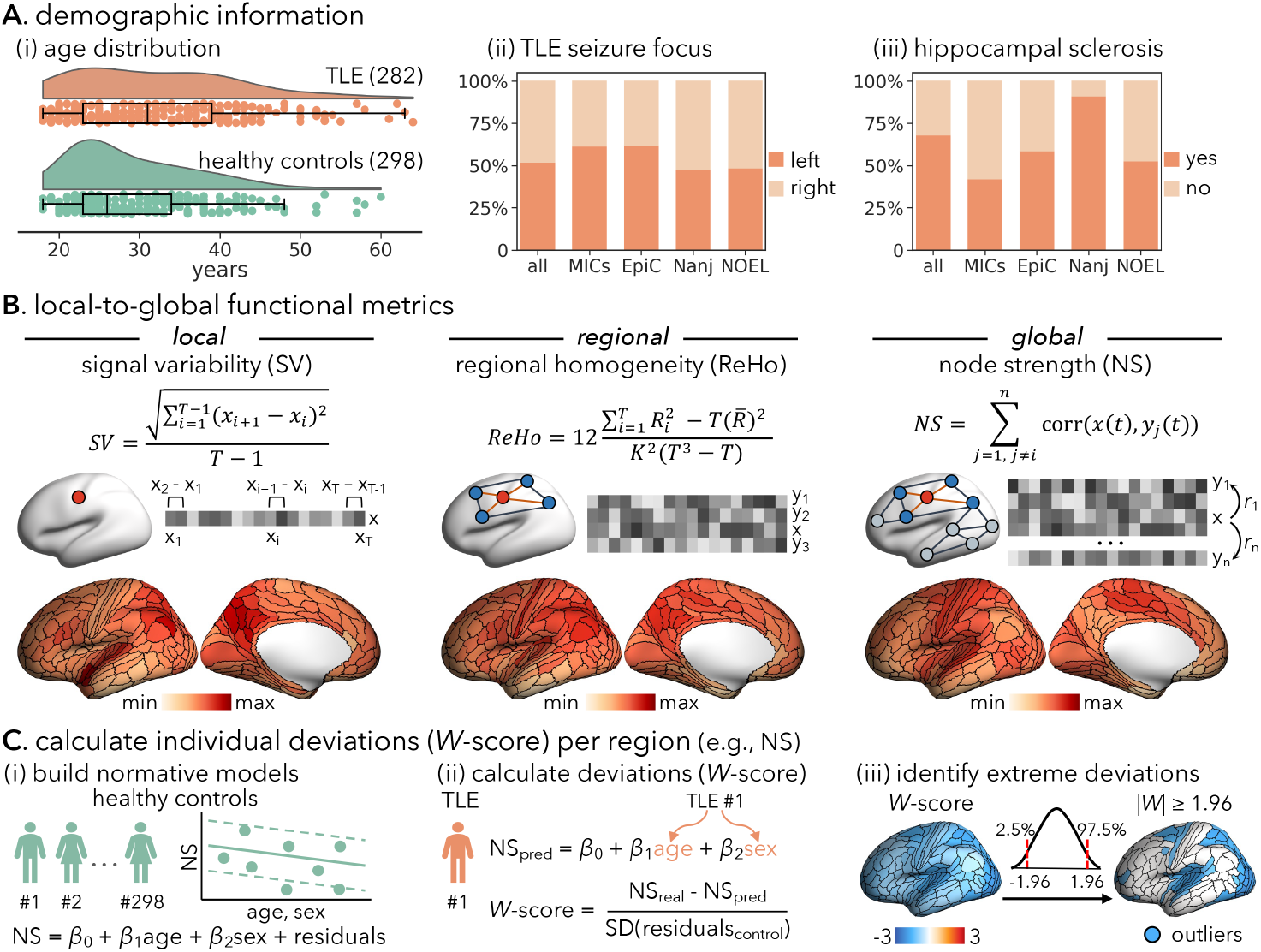
Overview of participants and analysis pipeline. **(A)** Demographic characteristics of the healthy control (*n* = 298) and TLE groups (*n* = 282). (*i*) Age distributions. Boxes represent the interquartile range (IQR), with the lower and upper boundaries corresponding to the 25th and 75th percentiles. Whiskers extend to the minimum and maximum values within 1.5×IQR from the 25th and 75th percentiles. Each dot represents an individual participant. (*ii*) Proportion of TLE patients with left-or right-sided seizure focus in each dataset. (*iii*) Proportion of TLE patients with or without ipsilateral hippocampal sclerosis in each dataset. **(B)** Overview of methodology for calculating functional metrics from resting-state functional MRI (rs-fMRI) time series in each brain region: signal temporal variability, regional homogeneity, and node strength. **(C)** Schematic of individual functional deviation (*W*-score) estimation, with an example here for node strength (NS). (*i*) Building normative models in healthy individuals. Age-and sex-related variations in each rs-fMRI metric are modeled using a linear regression model in the healthy control group, yielding beta maps for intercept (*β*0), age (*β*1), sex (*β*2), and standard deviation of residuals in each brain region. (*ii*) Estimating deviations in patients. Predicted value for a given patient’s age and sex (NSpred) is calculated as *β*0 + *β*1×age + *β*2×sex. *W*-scores are defined as the normalized deviation of the observed values from the corresponding predicted values. (*iii*) Identifying extreme deviations. Brain regions with *W*-scores exceeding ±1.96 (*i*.*e*., |*W*-scores| ≥ 1.96) were classified as showing extreme deviations, corresponding to the upper and lower 2.5% of the normative distribution.

### 2.2 Region-Specific Functional Deviations in TLE

Three metrics, chosen to index brain function at 3 different spatial scales (local, regional, and global), were calculated from processed rs-fMRI time series at the node level: signal temporal variability (SV), regional homogeneity (ReHo), and node strength (NS) (**Figure 1B**). Normative percentiles of variations in SV, ReHo, and NS were established using rs-fMRI data from healthy individuals. Deviations from the percentile curves were quantified with *W*-scores (**Figure 1C**).^20,32^ Each brain region for each patient was classified as either an extreme deviation (|*W*-scores| ≥ 1.96) or within the normative range, generating an individualized deviation map.^24-26^ Compared to healthy controls, individuals with TLE exhibited extensive yet variable alterations in SV, ReHo and NS. Across the cortex, the median proportion of extreme deviations in SV, ReHo, and NS in TLE group was 5.3% (range = 1.1%–12.4%), 7.4% (2.1%–16.0%), and 6.4% (2.1%–11.4%), respectively (**Figure 2A, S1**). Stratifying findings according to lobes, extreme deviations in SV and ReHo were most prevalent in the temporal, parietal and occipital lobes, whereas NS deviations were more marked in the parietal and insular cortex, with a higher prevalence ipsilateral to seizure focus. Next, aggregating the three metrics using a multivariate Mahalanobis distance revealed a diffuse spatial pattern of functional deviations across the cortex in TLE patients (median: 5.3% [1.8%–12.4%], **Figure 2B**), with the highest overlap observed in the ipsilateral mesial temporal lobe (11.4%; **Figure S2**). These findings suggest that although extensive functional changes are a hallmark of TLE (97% of TLE patients had extreme deviations in at least 10 brain regions), the specific pattern of affected regions varied markedly across individuals. We further confirmed the consistency of these findings across datasets by estimating TLE-related functional deviations in each dataset independently and correlating them with the main finding in **Figure 2B** (spatial similarity: *rho* = 0.24–0.60, *P*_spin_ < 0.001; **Figure S3**). Notably, the subgroup of TLE patients with hippocampal sclerosis (HS) showed more extreme deviations in the ipsilateral (*t* = 2.61, *P*_FDR_ = 0.024) and contralateral (*t* = 2.75, *P*_FDR_ = 0.024) insular cortices compared to those without HS (**Figure S4**).

**Figure 2:**
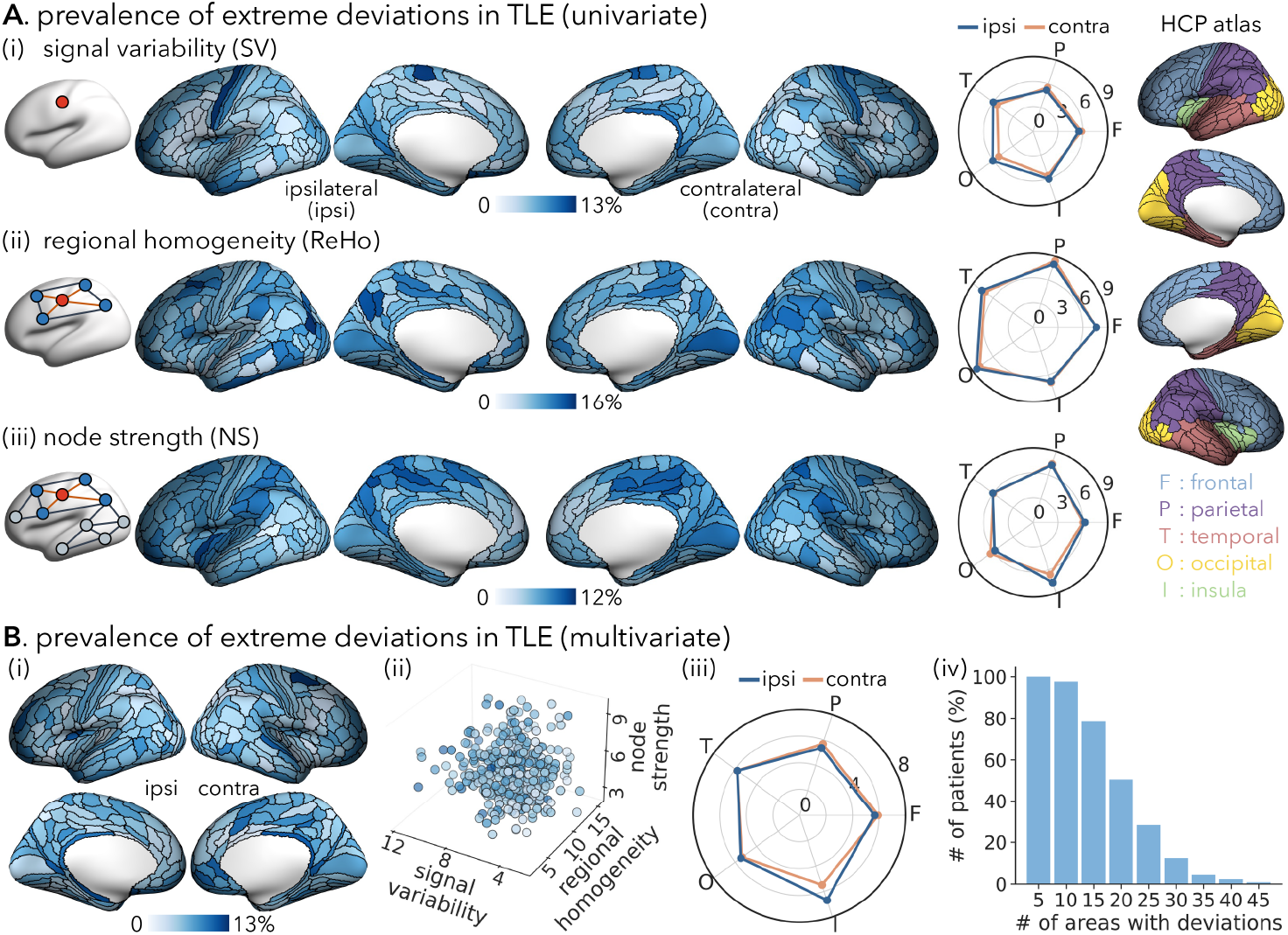
Region-specific prevalence of extreme functional deviations in TLE patients. **(A)** Proportion of TLE patients with extreme deviations (|*W*-score| ≥ 1.96) in each brain region for (*i*) signal variability (SV), (*ii*) regional homogeneity (ReHo), and (*iii*) node strength (NS). Spider plots show the mean proportion of extreme deviations in each cortical lobe defined on the HCPMMP1.0 atlas. **(B)** (*i*) Proportion of TLE patients with multivariate *W*-scores (aggregating SV, ReHo and NS) exceeding ±1.96. (*ii*) Region-specific deviation prevalence distribution. Each dot represents a brain region, with its position indicating the prevalence values of the three metrics and its color denoting the composite deviation prevalence. (*iii*) Mean proportion of extreme deviations per lobe. (*iv*) Distribution of the number of extreme deviations per patient.

### 2.3 Network Constraints on Regional Functional Deviations

We next explored the extent to which the spatial pattern of functional deviations was shaped by white matter network connectivity among brain regions. Specifically, we examined whether the structural connectivity profile of a given region *i* could predict the functional deviation scores of other regions to which region *i* was structurally coupled. This approach has previously been used to map epicentres of disease-related alteration patterns.^3,5,33^ For each brain area, we calculated the mean deviation prevalence value of its connected neighbors, weighted by white matter connectivity strength estimated by diffusion MRI streamlines tractography (**Figure 3A**). A positive correlation was observed between the node and mean neighbor deviation maps (*rho* = 0.27, *P*_spin_ < 0.001). We further identified disease epicenters, defined as brain regions wherein seed-based structural connectivity profiles spatially resembled the empirical functional deviation pattern in TLE (**Figure 3B**).^3^ The ipsilateral mesiotemporal lobe, bilateral medial prefrontal, and parietal cortices emerged as epicenters of functional alterations (*P*_spin_ < 0.05; **Figure 3C**).

**Figure 3:**
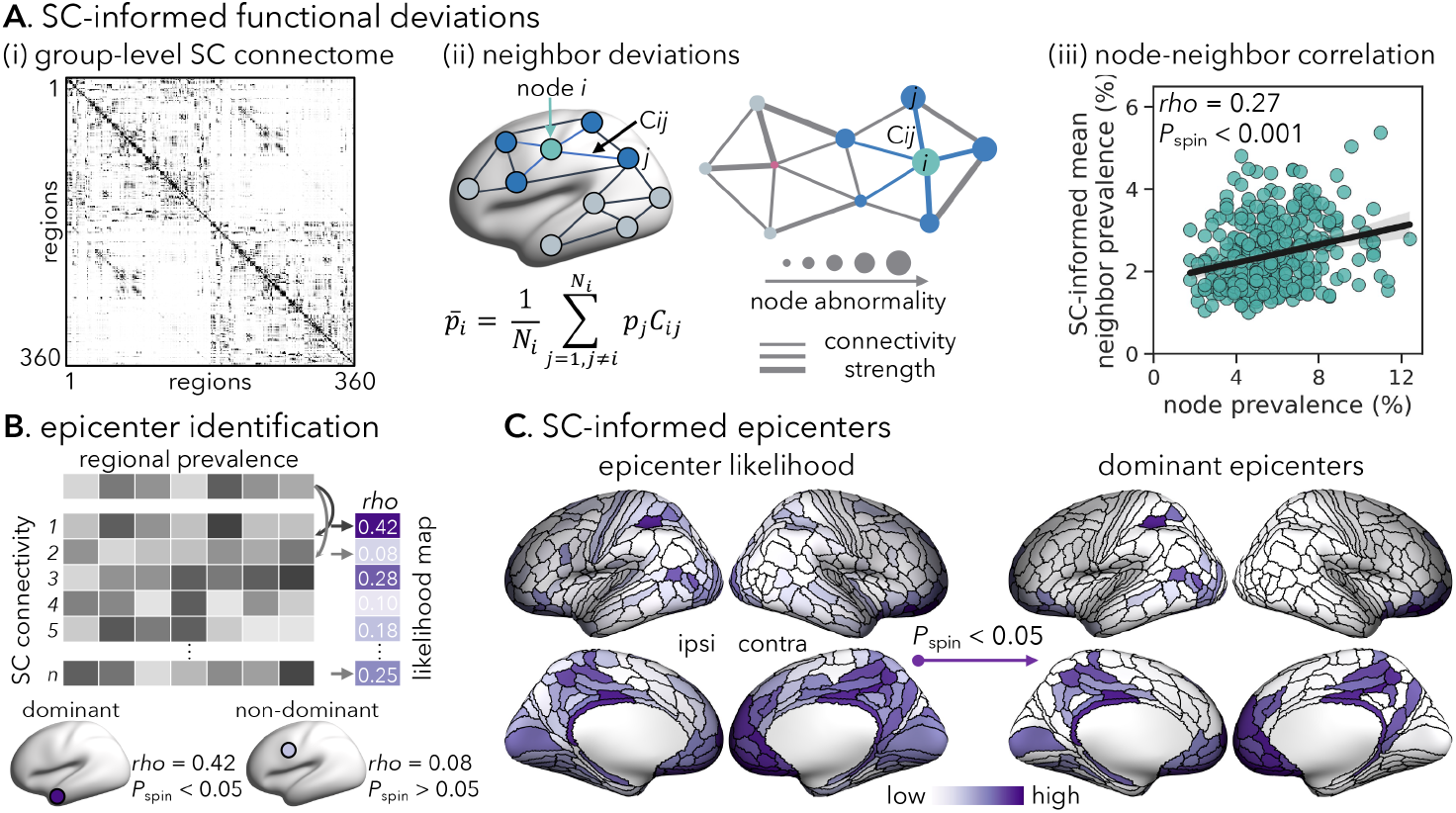
Network-based spreading of regional functional deviations. **(A)** (*i*) Group-level structural connectivity (SC) matrix from diffusion MRI of 100 unrelated healthy individuals. (*ii*) Schematic of functional deviations of a node (*pi*) and its neighbors 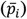. If the regional deviation depends on SC network organization, nodes connected to highly abnormal neighbors (*i*.*e*., high prevalence) will be more likely to be affected, whereas nodes connected to healthy neighbors (*i*.*e*., low prevalence) will be less likely to be affected. (*iii*) The functional deviation of a node (x-axis) is positively correlated with the mean deviation of neighbors (y-axis) to which they are structurally connected. Statistical significance (*i*.*e*., *P*spin) of the correlation coefficient is assessed using spin permutation tests (5,000 iterations). **(B)** Schematic of disease epicenter identification. A node whose SC profile spatially strongly relates with the TLE-related functional deviation map in **Figure 2B** is considered a disease “epicenter”. Epicenter likelihood is defined as the Spearman correlation coefficient between two brain maps. **(C)** SC-informed epicenter likelihood map. Statistical significance of the likelihood is determined using spin permutation tests (5,000 iterations and *P*spin < 0.05). ipsi = ipsilateral; contra = contralateral.

### 2.4 Associations with Structural Deviations

We next examined the association between functional and structural alterations in TLE, using markers of whole-brain structural compromise (*i*.*e*., CT, FA, MD) widely used in studies of this condition.^34,35^ Like functional metrics, individualized structural deviations were computed using T1-weighted and diffusion-weighted MRI features by mapping each patient’s data onto normative percentile charts of healthy individuals. Across the cortex, structural deviations were most marked in the temporal lobe (mean CT/FA/MD = 7.5%/14.7%/16.6%), followed by the insular cortex (8.2%/9.6%/10.6%) and, to a lesser extent, the parietal lobe (7.8%/7.6%/10.1%; **Figure 4A**). To investigate the relationship between structural and functional alterations, we fitted a multilinear regression model to predict cortex-wide functional deviation’s prevalence based on structural deviations’ prevalence. The model demonstrated a strong fit between empirical and predicted functional deviation patterns, above and beyond the effect of spatial autocorrelation (*rho* = 0.27, *P*_spin_ < 0.001; **Figure 4B**). Dominance analysis then showed that the mean diffusivity contributed the most to the model fit (56%), while cortical thickness and fractional anisotropy had modest contributions of 24% and 20%, respectively (**Figure 4C**).

**Figure 4:**
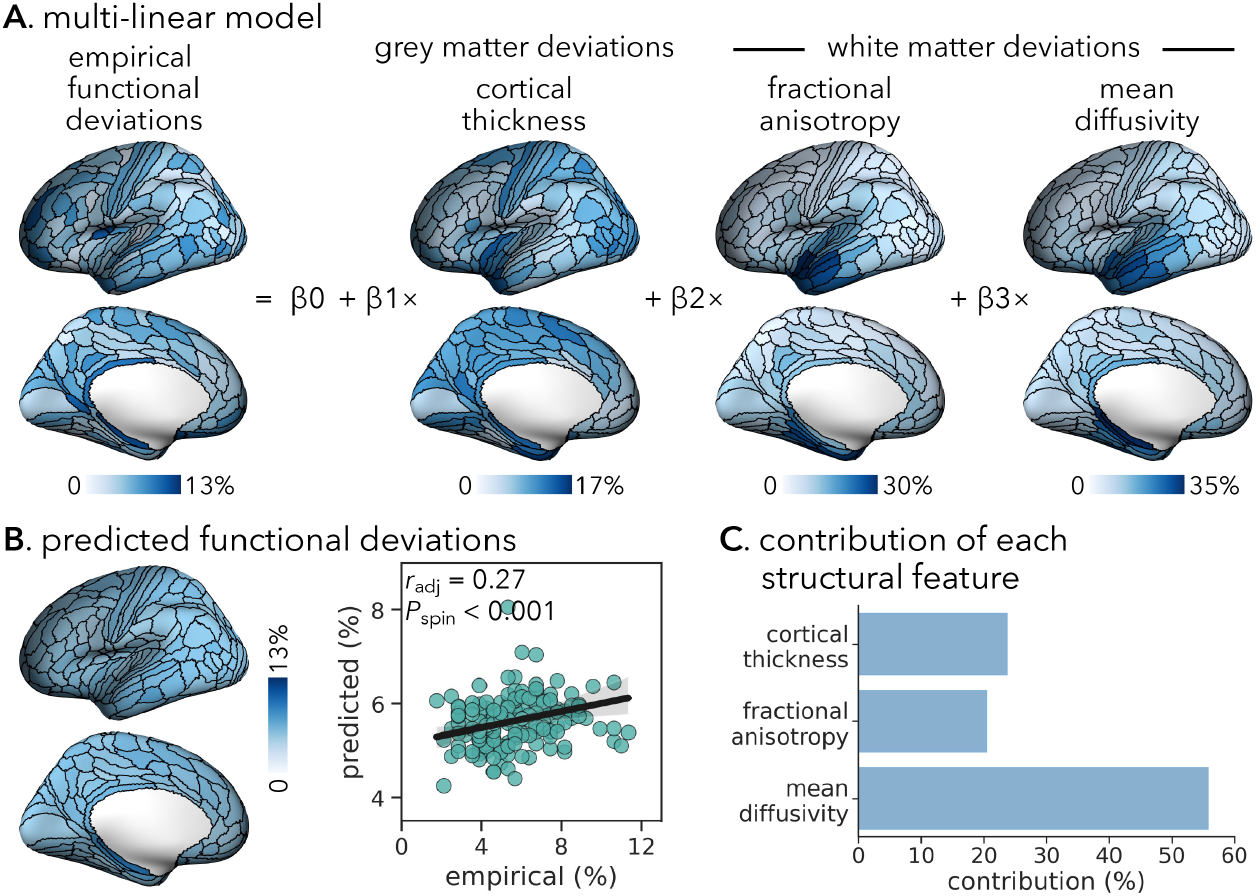
Spatial associations between brain structural and functional deviations. **(A)** A multilinear regression model is used to predict brain functional deviation prevalence from deviation prevalence maps of grey matter morphology (cortical thickness [CT]) and superficial white matter microstructure (fractional anisotropy [FA], mean diffusivity [MD]). **(B)** Spatial correlation between empirical (x-axis) and predicted (y-axis) functional deviation prevalence maps, with each dot representing a brain region. Statistical significance (*i*.*e*., *P*spin) of the correlation coefficient is determined using spin permutation tests with 5,000 iterations. **(C)** Dominance analysis quantifies the contribution of each feature (CT, FA and MD).

### 2.5 Clinical Utility of Functional Deviations

#### 2.5.1 TLE versus non-TLE Classification

Supervised machine learning models trained on individualized functional deviation metrics achieved an AUC of 0.76 (0.53–0.87) in discriminating TLE patients from non-TLE patients (*i*.*e*., FCD-related extratemporal focal epilepsy), which significantly exceeded chance levels (*P*_perm_ < 0.001; **Figure 5A**).

**Figure 5:**
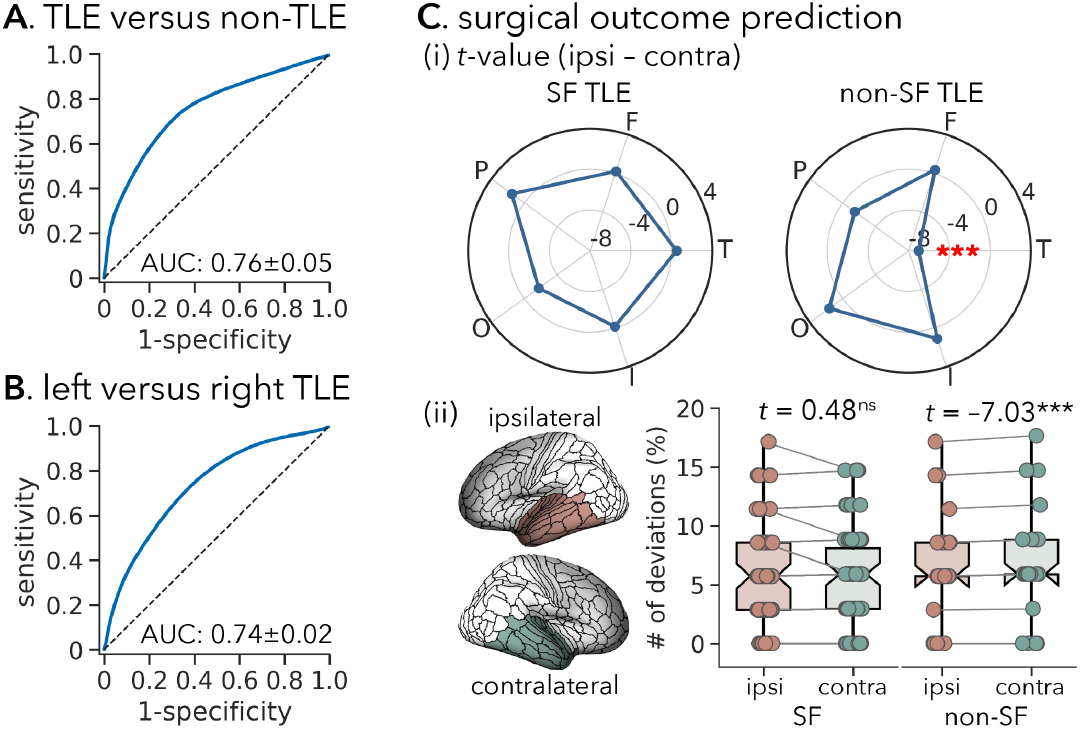
Clinical utility of individual functional deviations. (**A, B**) Receiver operating curves showing the accuracy of functional deviations in classifying TLE patients versus extratemporal FCD patients and left versus right TLE patients. (**C**) Interhemispheric differences in extreme deviations in seizure-free (SF) and non-seizure-free (non-SF) patients. (*i*) *T*-values showing the differences in the number of extreme deviations between the ipsilateral (ipsi) and contralateral (contra) hemispheres in SF and non-SF patients in each cortical lobe, respectively. Positive/negative *t*-values indicate more/fewer extreme deviations in the ipsilateral hemisphere compared to the contralateral hemisphere. (*ii*) Scatter plots showing the patient-specific proportions of extreme deviations in the ipsilateral (red) and contralateral (green) temporal lobes in SF and not-SF patients. T = temporal; F = frontal; P = parietal; O = occipital; I = insula. *** *P*FDR < 0.001.

#### 2.5.2 Seizure Focus Lateralization

Patients with left TLE could be distinguished from those with right TLE with an AUC of 0.74 (0.65– 0.80) based on individualized functional deviation scores (*P*_perm_ < 0.001; **Figure 5B**).

#### 2.5.3 Postsurgical Seizure Outcome Prediction

We examined whether inter-individual differences in preoperative functional deviations were associated with postsurgical seizure freedom in TLE patients who underwent resective surgery (*n* = 99). A positive spatial correlation was observed between the prevalence map of extreme deviations and the overlap map of surgical cavities (derived from pre-and post-surgical T1-weighted MRIs; mean ± SD = 39% ± 37%, range = 3%–100%) across regions (*rho* = 0.19, *P*_spin_ = 0.07; **Figure S5**), suggesting a higher likelihood of functional anomalies in to-be-resected brain regions. Moreover, in patients with postsurgical seizure recurrence, we observed greater numbers of extreme deviations in the contralateral temporal lobe compared to the ipsilateral hemisphere (*t* = –7.03, *P*_FDR_ < 0.001; **Figure 5C**). In contrast, no such a significant difference was found in seizure-free patients (ipsilateral versus contralateral, *t* = 0.48, *P*_FDR_ = 0.407). There were no significant interhemispheric differences in other lobes for either subgroup (ipsilateral versus contralateral, *P*_FDR_ > 0.084). The supervised machine learning model based on functional deviation scores discriminated seizure-free versus non-seizure-free patients with an AUC of 0.63 (*P*_perm_ = 0.023).

#### 2.5.4 Associations with Clinical Characteristics

A positive correlation was observed between disease duration and the number of extreme deviations in the TLE cohort (*r* = 0.09, *P* = 0.070), indicating more diffuse functional deviations in patients with long-standing TLE. No significant correlations were found between the number of extreme deviations and age at seizure onset or the number of antiseizure medications (*P* > 0.235). However, patients with a history of focal to bilateral tonic-clonic seizures exhibited significantly more deviations than those without focal to bilateral tonic-clonic seizures (*t* = 1.73, *P* = 0.042).

### 2.6 Sensitivity Analyses

Several sensitivity analyses confirmed the robustness of our main findings.

#### 2.6.1 Left and Right TLE

We recomputed the functional deviation prevalence at the group level separately in left and right TLE patients. Overall, findings were spatially similar (**Figure S6**); however, left TLE patients additionally had more extreme deviations in the temporal lobe, while right TLE patients showed more deviations in the parietal lobe.

#### 2.6.2 Head Motion and Global Mean Signal

The spatial distribution of functional deviations in TLE patients was virtually identical when controlling for individual head motion during rs-fMRI scans (median: 5.3% [1.8%–12.8%]; *rho* = 0.97, *P*_spin_ < 0.001) and global mean signal (median: 5.3% [1.8%–12.1%]; *rho* = 0.71, *P*_spin_ < 0.001; **Figure S7**).

## 3. Discussion

In this study, we quantified and visualized interindividual differences in patterns of cortical functional alterations in a large, multicentric dataset of individuals with TLE alongside with healthy controls and a disease control group with FCD-related extratemporal epilepsy. Although brain functional deviations were most prominent in the temporal lobe, there remained substantial variability across patients, indicating that TLE disrupts large-scale brain functional organization in a non-uniform manner even within its typical temporolimbic focus. The spatially distributed pattern of functional alterations nevertheless aligned with the brain’s structural connectome architecture, suggesting a pivotal role of white matter tracts as conduits for the propagation of functional disturbances.^5,34,36^ Paralimbic and medial default mode cortices emerged as epicenters of macroscale dysfunction, consistent with the presumed pathological substrate of TLE. These functional changes were primarily reflective of concurrent white matter microstructural compromise, followed by cortical thinning. Machine learning models informed by individual functional deviations successfully identified diagnostic categories and seizure focus laterality with medium-to-high accuracy and predicted postsurgical seizure outcome with moderate accuracy. Collectively, our findings show that functional alterations in TLE patients are common yet spatially heterogeneous, demonstrating the promise of patient-specific brain deviation mapping for clinical diagnosis and prognosis.

Studying multicentric rs-fMRI data from four independent datasets, we employed a normative modeling approach to quantify individual patient deviations in brain function and generate region-specific prevalence maps of extreme deviations in patients with TLE. Compared to healthy controls, TLE patients presented with extensive alterations in SV, ReHo and NS, commonly affecting temporolimbic and parietal cortices, albeit with relatively low proportions of significant deviations across patients (up to 16%). Despite this variability, 97% of TLE patients had extreme deviations in at least 10 brain areas. In particular, the most evident deviations were seen in the ipsilateral mesial temporal structures. These are areas known to be implicated in the pathogenesis of TLE and are responsible for its clinical manifestation.^37,38^ While alterations in SV, ReHo and NS are indeed core pathopsychological features evident in most individuals with TLE,^4,11,34,39-41^ our findings suggest that the specific cortical regions affected by these abnormalities differ markedly between patients. TLE impacts individuals in such a heterogenous manner so that group-consensus maps of TLE pathology derived from classic disease-control inferential paradigms may not fully capture the full spectrum of individual disease manifestations.^15,16,42^ Notably, in TLE patients with HS, the overall pattern of functional deviations remained largely similar; however these patients exhibited greater deviations in the ipsilateral hemisphere (particularly the insula), compared to those without HS. These findings are in line with recent MRI-based research describing greater changes in excitation/inhibition balance, hippocampal functional connectivity, and network controllability in HS-TLE patients.^5,43,44^ Moreover, they confirm previous *postmortem* work that has reported marked extramesial cortical pathology in HS-TLE patients, often also extending towards temporopolar, frontopolar, and orbitofrontal regions.^45^

The human brain is an intricate network of functionally specialized brain regions. Structural connections between these regions allows for the coordination of functional interactions and the transport of trophic signals.^46-49^ They can also act as conduits for the spread of pathology, such that focal perturbations can propagate to impact distributed systems.^33,50,51^ Our findings align with this theory, demonstrating that functional deviations in TLE are more tightly coupled between regions with strong structural connectivity. While our study is cross-sectional in nature, we posit that connectome architecture exerts a strong influence on the spread of cortical functional damage in TLE. Such functional alterations may arise from a transsynaptic network process,^50,51^ in which dysregulations of the pathophysiological core may trigger atypical activity in adjacent brain structures that, over time, may result in widespread brain dysfunction affecting multiple systems. Prior studies have shown that patterns of tissue volume loss in TLE follow the brain’s structural network embedding, with brain atrophy originating in the mesiotemporal cortices at early stages of the disease, subsequently affecting extratemporal tissues, as well as structurally connected regions in distant neural territories.^3,7,8,52^ Our study extends this work by showing that the spatial pattern of functional alterations is also shaped by the underlying structural connectome, reinforcing the fundamental role of global network architecture in shaping cortical abnormalities in TLE. In addition, several studies have demonstrated that brain regions are targeted non-randomly by disease; those that are highly connected and potentially important for communication tend to be disproportionately affected.^53^ Indeed, our epicenter mapping analysis identified key areas implicated in functional disruption, primarily in regions often considered to belong to transmodal association cortex, such as the medial temporal, prefrontal, and parietal regions.^3,5^ The association cortex is a core component of the brain’s putative rich-club—densely interconnected regions that are thought to facilitate global signal integration and information broadcasting.^54^ As network disconnection of the association cortex has been well documented in TLE,^4,55^ we build upon prior work by showing that medial prefrontal and parietal regions are both affected and likely relate to distributed network-level propagation of functional imbalances. Altogether, our analysis looking beyond regional changes and considering large-scale alterations in brain organization highlights the interconnected nature of TLE pathology, emphasizing that macroscale functional disturbances are strongly shaped by the brain’s structural architecture, and motivates efforts to explore the potential of network-level interventions as therapeutic strategies.^3,56^

Our multimodal MRI framework enables us to investigate the relationship between functional deviations and structural compromise in the same cohort, with a particular focus on patient-level variability. Notably, we found the most marked deviations in TLE in the ipsilateral temporal lobe, affecting grey matter morphology (CT) and white matter integrity (FA and MD), with varying degrees of severity. Through a multilinear regression model, we revealed a close fit between observed functional deviation maps and maps predicted on the basis of structural deviations, suggesting that structural damage partially accounts for functional alterations in TLE. Unlike earlier studies that primarily explored group-averaged patterns of brain changes,^4,36^ this study emphasizes coupled interindividual variability in both structural and functional alterations. This individualized approach is highly relevant in TLE, where neuroanatomical changes have shown to vary heavily between patients in both mesiotemporal and neocortical systems.^6,38,57^ Notably, additional dominance analysis revealed that MD contributed most to the predictive relationship, underscoring a critical role of white matter integrity in brain function and dysfunction.^4,11^ White matter pathways serve as the structural backbone for effective communication between brain regions, facilitating the transmission of neural signals, synchronization of activity, and coordination of functional networks.^58,59^ Accordingly, white matter microstructural impairment, as observed in TLE, may disrupt local brain activation or long-range functional connectivity, contributing to the breakdown of large-scale networks such as the default mode and limbic networks, which are commonly implicated in TLE.^4^ Our results suggest that cortical functional deviations in TLE are not merely reflective of group-level structural changes across patients, but are closely linked to each patient’s unique pattern of white matter damage. This patient-specific perspective highlights the importance of personalized assessments in capturing diverse structural and functional profiles of TLE, as well as in understanding pathophysiological mechanisms driving disease progression in each patient. Here, we provide a framework for future studies to track the evolution of individual structural and functional changes over time, providing new opportunities for disease monitoring. Capturing each patient’s unique pattern can therefore deepen our understanding of the structure-function interplay underlying TLE at the individual patient level.

Normative modeling shows great promise for personalized clinical decision-making in TLE. Leveraging supervised machine learning technique, we provided evidence that individualized maps of functional deviations distinguished TLE patients from those with FCD-related extratemporal epilepsy and identified TLE seizure onset zone laterality with a high accuracy. The differentiation between left and right TLE patients was primarily driven by functional deviations in the temporal and parietal cortices ipsilateral to seizure focus. Specifically, left TLE exhibited greater deviations in the temporal lobe, which is in alignment with known decreases in functional connectivity of the lateral temporal lobe.^60^ It remains to be determined whether these observations overall relate to differences in clinical and cognitive phenotypes, and whether they suggest variable pathophysiological substrates of left versus right TLE.^61,62^ The approach developed here may inform and ultimately enhance the presurgical evaluation of pharmaco-resistant TLE using non-invasive MRI techniques, which has traditionally primarily focused on structural MRI.^63-65^ Future translational research should explore the combined utility of structural and functional MRI data to further improve clinically relevant diagnostics.

In the subgroup of TLE patients who underwent temporal lobe resections, we observed a significant, yet modest association between preoperative functional alterations and postoperative seizure outcome. Specifically, patients with postsurgical seizure recurrence exhibited greater functional deviations in the temporal lobe contralateral to seizure focus. That is, more distributed abnormalities extending to the contralateral hemisphere are, therefore, tied to unfavorable prognosis. This is consistent with clinical intuition,^66,67^ as well as previous findings indicating that aberrant functional connectivity of the contralateral temporal lobe reflects poorer postsurgical seizure outcome prospects.^68,69^ Persistent aberrant activity in non-surgically resected brain regions increases the likelihood of seizure recurrence.^5,68^ Our finding aligns with this consensus, and with the notion that “temporal plus” epilepsy may contribute to temporal lobe surgery failures.^70^ In such cases, the presence of contralateral functional anomalies may contribute to ongoing epileptogenic activity after a unilateral resection. Therefore, preoperative identification of structural and functional signatures using non-invasive methods may help to determine and refining treatment strategies. In particular, neuromodulation technique is increasingly recognized as an effective alternative for TLE patients who continue to suffer from seizures following resective surgery.^71,72^ The individualized functional deviation mapping framework proposed in this study could aid in pinpointing precise targets, paving the way for patient-specific treatments.

## 4. METHODS

### 4.1 Participants

In the TLE cohort, no patients had mass lesions (tumor, vascular malformations, or malformations of cortical development) and none had a history of traumatic brain injury or encephalitis. Among the included patients, 75.9% of TLE patients (*n* = 214) and 88.9% of FCD patients (*n* = 40) had drugresistant seizures. For TLE patients, the mean ± SD age at seizure onset was 17.8 ± 10.7 years [0.3– 60 years] with a mean ± SD disease duration of 14.4 ± 11.0 years [0.8–49 years]. For FCD patients, the mean ± SD age at seizure onset was 11.5 ± 6.9 years [0.3–27 years], with a mean ± SD disease duration of 15.9 ± 10.3 years [0.5–48 years]. TLE diagnosis and lateralization of the seizure focus as left (*n* = 147; mean age = 31.7 ± 10.6 years [18–64 years]) or right TLE (*n* = 135; mean age = 32.9 ± 9.3 years [18–63 years]) were determined following the ILAE criteria through comprehensive evaluations,^73^ including clinical history, seizure semiology, video-EEG recordings, clinical MRI, and/or neuropsychological assessment. In our cohort of TLE patients, 68.1% (*n* = 192) had ipsilateral hippocampal sclerosis (HS) (**Figure 1A**), confirmed by radiological MRI reports, quantitative analyses of T1-weighted and FLAIR images, and histological assessments;^74-76^ 6.4% (*n* = 18) had ipsilateral hippocampal gliosis; and 25.5% (*n* =72) were MRI-negative. Among the 99 TLE patients who underwent temporal lobe resections, 74 (75%) achieved seizure freedom (Engel class I) and 25 (25%) continued to experience seizures (Engel class II–IV) at a mean follow-up period of 44 ± 34 months. The Institutional Ethics Committee at each institution approved the study procedures. All participants provided written informed consent in accordance with the Declaration of Helsinki.

### 4.2 MRI Processing

Multimodal MRI data, including T1-weighted, diffusion MRI, and rs-fMRI, were acquired with 3.0-Tesla scanners (*MICA-MICs*: Siemens Prisma; *EpiC*: Philips Achieva; *Nanj* and *NOEL*: Siemens Trio) in all individuals (prior to surgery for patients). MRI data of the four datasets were processed uniformly using *micapipe* (version 0.2.3; http://micapipe.readthedocs.io) and mapped to cortical surface points (henceforth, vertices).^77^ We extracted the rs-fMRI BOLD time series, cortical grey matter morphology, and white matter microstructural features for each participant using the same set of 360 cortical parcels (HCP-MMP1.0 atlas).^78^ Full details on MRI data acquisition and processing are provided in the **Supplementary Materials**.

### 4.3 Resting-State fMRI Metrics

Signal variability (SV), a local metric of neural signal fluctuations over time, was defined as the root mean squared successive difference between time points of rs-fMRI time series.^79^

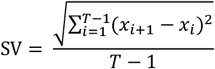

where *x*_*i*_(*t*) is the rs-fMRI signal at the *i*-th time point for a node *x*, with time series length *T*: *x* = (*x*_1_, *x*_2_,…, *x*_*T*_). Higher/lower SV indicates stronger/weaker temporal fluctuations. Although SV has been widely used to index local neural function in both healthy and clinical populations,^79-81^ no systematic assessment has yet been conducted in epilepsy.

Regional homogeneity (ReHo), as an index of regional brain function, was computed as the temporal coherence of unsmoothed rs-fMRI time series within a given vertex’s nearest regional neighbors.^82^

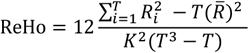

where *R*_*i*_ is the ranks of time series *x*_*i*_(*t*), and 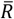is the mean rank across all *K* neighbors and time points. Higher/lower ReHo represents stronger/weaker local functional homogeneity. ReHo has been widely applied to assess regional subnetwork coherence in both healthy individuals and clinical populations, including individuals with TLE.^40,82^

To assess the global connectivity of a given vertex, node strength (NS) was defined as the sum of the weighted functional connectivity strengths, calculated as the Pearson correlation coefficient between rs-fMRI time series of a given brain region and all other regions.^83^

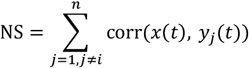

where *y*_j_(*t*) represents the rs-fMRI time series for a node, excluding node *i*, and *n* = 360 is the total number of nodes. A higher NS indicates greater similarity in intrinsic neural activity between regions, whereas a lower NS reflects greater inter-regional dissimilarity. In both the study of healthy and neurological patient populations,^28,84^ NS has been extensively used to chart globally interconnected hub regions and their disruption.

### 4.4 Individualized *W*-score Maps

To account for potential variations between scanners/datasets, we applied ComBat to harmonize each rs-fMRI metric independently,^85^ with each dataset treated as a batch while retaining variances of interest (age, sex, and diagnosis). ComBat leverages multivariate linear mixed-effects regression and empirical Bayes to correct batch effects.^86^ The harmonized data correcting for inter-dataset variations were then used to generate *W*-score maps.

For each metric and each brain region, a *W*-score was calculated using the healthy control group as a reference.^20,32^ Analogous to the *Z*-score^23^ with a mean value of 0 and a standard deviation of 1 relative to healthy controls, the *W*-score reflects normalized deviation of the observed value from the normative range for healthy adults of the same age and sex. Here, age-and sex-related changes in each rs-fMRI metric were estimated in the healthy control group through a linear regression model (*β*_0_ + *β*_1_ × age + *β*_2_ × sex + residuals) (**Figure 1C**), yielding beta maps for intercept value (*β*_0_), age-(*β*_1_) and sex-related (*β*_2_) coefficients, as well as individual maps of residuals for each brain region. Patient-specific *W*-scores were estimated using the formula:^20,32^

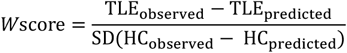

where the value predicted for a given patient’s age and sex (TLE_predicted_) was calculated as: *β*_0_ + *β*_1_ × age + *β*_2_ × sex, and SD represented the standard deviation of model residuals in healthy controls. To this end, deviations from normative distributions of SV, ReHo and NS were generated for each brain region and each patient. Positive and negative *W*-scores indicate deviations above and below the normative ranges, respectively. Brain regions with *W*-scores exceeding ±1.96 were classified as showing extreme deviations (above and below the 2.5th percentiles)^24-26^. The proportion of extreme deviations (*i*.*e*., group-level deviation rate) was computed by counting patients with |*W*-scores| ≥ 1.96 and dividing by the total patient count in each brain region and was stratified across five lobes (*i*.*e*., frontal, parietal, temporal, occipital, and insular). Finally, to aggregate the three rs-fMRI metrics, we built a composite score for each individual, defined as the Mahalanobis distance of the joint distributions,^34^ and used the map in our multivariate *W*-score calculations.

### 4.5 Neighborhood Deviation Estimates

A group-averaged structural connectivity matrix, derived from an independent sample of 100 healthy adults^87^ was used to define the neighbors of a given brain region. The collective deviation prevalence value 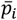 for neighbors of the *i*-th brain region was estimated as the average weighted prevalence value of all brain regions structurally connected to region *i* (*i*.*e*., regions with no structural connectivity to region *i* were excluded):^5,88^

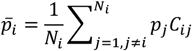

where *N*_*i*_ was the number of connected neighbors of region *i* (*i*.*e*., node degree), *p*_j_ was the deviation prevalence value observed in the *j*-th neighbor of region *i*, and *C*_*ij*_ was the connectivity strength between *i* and *j*. Under this model, neighbors of region *i* with a more strongly weighted connection made stronger contributions to estimating region *i*’s neighborhood brain function change. Spearman correlation was used to assess the relationship between node and mean neighbor prevalence values. Statistical significance of correlation coefficients between cortical maps was determined using spatial autocorrelation-preserving spin tests.^89^ This procedure was repeated 5,000 times, generating a null distribution of brain maps with preserved spatial autocorrelation. The *P*-value (*i*.*e*., *P*_spin_) was calculated as the fraction of correlation coefficients in null models that exceeded the empirical correlation coefficient.

### 4.6 Disease Epicenters Mapping

Disease epicenters were identified by spatially relating each brain region’s healthy structural connectivity profile (from the same independent dataset as above^87^) to group-level functional deviation prevalence map in TLE.^3^ This approach was repeated systematically for each brain region, and the significance of spatial similarity was determined using spin permutation tests with 5,000 iterations. Brain regions with significant spatial correlations (*P*_spin_ < 0.05) were ranked in descending order according to the epicenter likelihood (*i*.*e*., Spearman correlation coefficients), with highly ranked regions representing potential disease epicenters.

### 4.7 Associations with Structural Deviations

Using the same framework as for rs-fMRI metrics, we generated individual *W*-score maps for cortical morphology (cortical thickness [CT]) and superficial white matter microstructure (fractional anisotropy [FA], mean diffusivity [MD]). Group-level prevalence maps for extreme deviations (|*W*-scores| ≥ 1.96) were created for each structural metric.

A multilinear regression model was employed to assess the associations between structural and functional deviations across the cortex (**Figure 4A**):

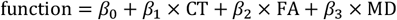

where the dependent variable was the group-level functional deviation prevalence map, and the independent variables were structural deviation prevalence maps. The intercept *β*_0_ and regression coefficients (*β*_1_, *β*_2_, *β*_3_) were optimized to maximize the spatial correlation between the empirical and predicted functional deviation prevalence maps. Model fit was quantified using the adjusted-*R*^2^ (coefficient of determination).

Dominance analysis was conducted to determine the relative contributions of structural metrics to the prediction of brain function deviation prevalence.^90^ Briefly, dominance analysis estimated the relative importance of each independent variable by constructing all possible combinations of variables and refitting the multilinear regression model for every combination. The contribution of each variable was quantified as the increase in explained variance (*i*.*e*., gain in adjusted-*R*^2^) when that variable was added to the model. Dominance scores were normalized by total model fit, enabling comparisons across models.

### 4.8 Clinical Utility of Functional Deviations

A supervised machine learning algorithm, implemented in LIBSVM,^91^ was used to evaluate the utility of individual functional deviation scores in classifying diagnoses, seizure onset zone laterality, and postsurgical outcome. Model training and testing used a nested 5-fold cross-validation with 1,000 iterations. Specifically, in each outer iteration, 4 folds were used for training and the remaining fold for testing until all folds had served as training and testing sets. An inner 5-fold cross-validation loop was applied to the training set to optimize hyperparameters (*c, g*) through grid search. Once the hyperparameters were determined, models were re-trained on the entire training set. Model performance was evaluated on the testing set using the area under the receiver operating characteristic curve (AUC). Statistical significance was determined using 1,000 permutation tests with randomly shuffled participant labels, with the *P*-value (*i*.*e*., *P*_perm_) calculated as the fraction of AUC values in null models that exceeded the empirical AUC.

#### 4.8.1 TLE versus non-TLE Classification

To assess the utility of functional deviation scores in identifying diagnostic categories, we trained an SVM model to distinguish TLE patients (*n* = 129) from extratemporal FCD patients (*n* = 45) in two datasets (*MICs* and *NOEL*) that had both cohorts available. Stratified splitting was employed to ensure consistent TLE/FCD ratios across all data splits.

#### 4.8.2 Seizure Focus Zones Lateralization

To evaluate the effectiveness of functional deviation scores in identifying seizure onset zone laterality, we trained a SVM model to classify left TLE patients (*n* = 147) versus right TLE patients (*n* = 135) using stratified splitting to preserve the left/right TLE ratio across all splits.

#### 4.8.3 Postsurgical Seizure Outcome Prediction

Postsurgical seizure outcome data were available for 99 TLE patients from 3 datasets (*MICs, NOEL*, and *Nanj*) who underwent temporal lobe resections: 74 (75%) were seizure-free (Engel class I), 25 (25%) were not seizure-fee (Engel class II–IV). Postsurgical MRI scans were available in 35 patients, enabling the segmentation of patient-specific surgical cavity mask from pre-and post-operative T1w MRIs. We first examined the spatial correspondence between the pattern of functional deviations and the pattern of between-patient overlap in surgical cavities, defined as the proportion of patients with resection in each region (out of 35 patients); next, we compared the proportion of extreme deviations (*i*.*e*., number of deviations divided by the total number of regions in each lobe) between the ipsilateral and contralateral hemispheres in each lobe for each TLE subgroup (seizure-free and non-seizure-free) using paired *t*-tests (*P*_FDR_ < 0.05); also, we trained an SVM model to classify seizure-free versus non-seizure-free TLE patients based on individual functional deviation scores using stratified splitting.

#### 4.8.4 Associations with Clinical Characteristics

Associations between the number of extreme deviations and disease course parameters (age at seizure onset, disease duration, and number of antiseizure medications) were evaluated in TLE patients using bivariate correlations. In addition, we split TLE patients according to the presence or absence of focal to bilateral tonic–clonic seizures and compared the number of extreme deviations between the two subgroups using two-sample *t*-tests.

### 4.9 Sensitivity Analyses

#### 4.9.1 Left and Right TLE

As the laterality of seizure focus may differentially affect the distribution of brain functional anomalies, we repeated calculating the group-level prevalence of extreme functional deviations in left (*n* = 147) and right (*n* = 135) TLE subgroups independently.

#### 4.9.2 Head Motion and Global Mean Signal

To assess the effects of head motion and global signal regression (GSR) on rs-fMRI data, we repeated the *W*-score analyses while additionally controlling for individual head motion (*i*.*e*., mean framewise displacement) during rs-fMRI scans, or using GSR-preprocessed data.

## Supporting information

Supplemental Methods and Results

## Acknowledgements

K.X. is funded by the China Scholarship Council (CSC, 202006070175) and the Healthy Brains and Healthy Lives (HBHL) Doctoral Fellowship. J.R. is funded by the Canadian Institutes of Health Research (CIHR). E.S., A.N., J.C. and R.R.C. are funded by the Fonds de Recherche du Québec - Santé (FRQ-S). S.L. is funded by the Centre de Recherche du CHUS. Lo.C. acknowledges support from a Berkeley Fellowship. L.C. is funded by the Consejo Nacional de Ciencia y Tecnología (CONACYT) (181508, 1782, FC218-2023) and Dirección General de Asuntos del Personal Académico (DGAPA) - UNAM (IB201712, IG200117, IN204720, IN213423). B.C.B. acknowledges research support from the National Science and Engineering Research Council of Canada (NSERC RGPIN-2025-05932), CIHR (FDN-154298, PJT-174995, PJT-191853), SickKids Foundation (NI17-039), Helmholtz International BigBrain Analytics and Learning Laboratory (HIBALL), HBHL, Brain Canada Foundation, FRQS, the Tier-2 Canada Research Chairs Program, and the Centre for Excellence in Epilepsy at the Neuro (CEEN).

## Data Availability

The *MICA-MICs* dataset is available at the Open Science Framework (https://osf.io/j532r/) and the Canadian Open Neuroscience Platform (https://portal.conp.ca/). The *EpiC* dataset is openly available at OpenNeuro (data set ds004469, https://openneuro.org/datasets/ds004469/versions/1.1.4). The *Nanj* and *NOEL* datasets will be made available upon reasonable request.

## References

1. Duncan JS, Winston GP, Koepp MJ, et al. Brain imaging in the assessment for epilepsy surgery. Lancet Neurol. 2016; 15:420–433.

2. Whelan CD, Altmann A, Botia JA, et al. Structural brain abnormalities in the common epilepsies assessed in a worldwide ENIGMA study. Brain. 2018; 141:391–408.

3. Larivière S, Rodriguez-Cruces R, Royer J, et al. Network-based atrophy modeling in the common epilepsies: A worldwide ENIGMA study. Sci Adv. 2020; 6:eabc6457.

4. Xie K, Royer J, Larivière S, et al. Atypical connectome topography and signal flow in temporal lobe epilepsy. Prog Neurobiol. 2024; 236:102604.

5. Xie K, Royer J, Rodriguez-Cruces R, et al. Temporal lobe epilepsy perturbs the brain-wide excitation-inhibition balance: associations with microcircuit organization, clinical parameters, and cognitive dysfunction. Adv Sci. 2025; 12:e2406835.

6. Lee HM, Fadaie F, Gill R, et al. Decomposing MRI phenotypic heterogeneity in epilepsy: a step towards personalized classification. Brain. 2021; 145:897–908.

7. Lee HM, Fadaie F, Gill RS, et al. MRI-derived modeling of disease progression patterns in patients with temporal lobe epilepsy. Neurology. 2024; 103:e209524.

8. Jiang Y, Li W, Li J, et al. Identification of four biotypes in temporal lobe epilepsy via machine learning on brain images. Nat Commun. 2024; 15:2221.

9. Hermann BP, Struck AF, Busch RM, et al. Neurobehavioural comorbidities of epilepsy: towards a network-based precision taxonomy. Nat Rev Neurol. 2021; 17:731–746.

10. Chen J, Alexander N, Raúl R-C, et al. A worldwide ENIGMA study on epilepsy-related gray and white matter compromise across the adult lifespan. bioRxiv. 2024;

11. Xie K, Royer J, Lariviere S, et al. Atypical intrinsic neural timescales in temporal lobe epilepsy. Epilepsia. 2023; 64:998–1011.

12. Reyes A, Kaestner E, Bahrami N, et al. Cognitive phenotypes in temporal lobe epilepsy are associated with distinct patterns of white matter network abnormalities. Neurology. 2019; 92:e1957–e1968.

13. Reyes A, Kaestner E, Ferguson L, et al. Cognitive phenotypes in temporal lobe epilepsy utilizing data-and clinically driven approaches: Moving toward a new taxonomy. Epilepsia. 2020; 61:1211–1220.

14. Rodriguez-Cruces R, Bernhardt BC, Concha L. Multidimensional associations between cognition and connectome organization in temporal lobe epilepsy. NeuroImage. 2020; 213:116706.

15. Marquand AF, Rezek I, Buitelaar J, et al. Understanding heterogeneity in clinical cohorts using normative models: Beyond case-control studies. Biol Psychiatry. 2016; 80:552–561.

16. Marquand AF, Kia SM, Zabihi M, et al. Conceptualizing mental disorders as deviations from normative functioning. Mol Psychiatry. 2019; 24:1415–1424.

17. Rutherford S, Barkema P, Tso IF, et al. Evidence for embracing normative modeling. eLife. 2023; 12:e85082.

18. Segal A, Tiego J, Parkes L, et al. Embracing variability in the search for biological mechanisms of psychiatric illness. Trends Cogn Sci. 2025; 29:85–99.

19. Mito R, Cole J, Genc S, et al. Towards precision MRI biomarkers in epilepsy with normative modelling. Brain. 2025; awaf090.

20. Renaud La J, Audrey P, Louisa B, et al. Region-specific hierarchy between atrophy, hypometabolism, and β-amyloid (Aβ) load in Alzheimer’s disease dementia. J Neurosci. 2012; 32:16265–16273.

21. Rutherford S, Kia SM, Wolfers T, et al. The normative modeling framework for computational psychiatry. Nat Protoc. 2022; 17:1711–1734.

22. Bethlehem RAI, Seidlitz J, White SR, et al. Brain charts for the human lifespan. Nature. 2022; 604:525–533.

23. Doering S, McCullough A, Gordon BA, et al. Deconstructing pathological tau by biological process in early stages of Alzheimer disease: a method for quantifying tau spatial spread in neuroimaging. eBioMedicine. 2024; 103:105080.

24. Bethlehem RAI, Seidlitz J, Romero-Garcia R, et al. A normative modelling approach reveals age-atypical cortical thickness in a subgroup of males with autism spectrum disorder. Commun Biol. 2020; 3:486.

25. Lv J, Di Biase M, Cash RFH, et al. Individual deviations from normative models of brain structure in a large cross-sectional schizophrenia cohort. Mol Psychiatry. 2020; 26:3512–3523.

26. Shao J, Qin J, Wang H, et al. Capturing the individual deviations from normative models of brain structure for depression diagnosis and treatment. Biol Psychiatry. 2024; 95:403–413.

27. Serena V, Seyed Mostafa K, Keir XXY, et al. Revealing individual neuroanatomical heterogeneity in Alzheimer disease using neuroanatomical normative modeling. Neurology. 2023; 100:e2442.

28. Sun X, Sun J, Lu X, et al. Mapping neurophysiological subtypes of major depressive disorder using normative models of the functional connectome. Biol Psychiatry. 2023; 94:936–947.

29. Bedford SA, Lai M-C, Lombardo MV, et al. Brain-charting autism and attention deficit hyperactivity disorder reveals distinct and overlapping neurobiology. Biol Psychiatry. 2025; 97:517–530.

30. Royer J, Rodriguez-Cruces R, Tavakol S, et al. An open MRI dataset for multiscale neuroscience. Sci Data. 2022; 9:569.

31. Weng Y, Lariviere S, Caciagli L, et al. Macroscale and microcircuit dissociation of focal and generalized human epilepsies. Commun Biol. 2020; 3:244.

32. Tremblay C, Abbasi N, Zeighami Y, et al. Sex effects on brain structure in de novo Parkinson’s disease: a multimodal neuroimaging study. Brain. 2020; 143:3052–3066.

33. Shafiei G, Markello RD, Makowski C, et al. Spatial patterning of tissue volume loss in schizophrenia reflects brain network architecture. Biol Psychiatry. 2020; 87:727–735.

34. Liu M, Bernhardt BC, Hong SJ, et al. The superficial white matter in temporal lobe epilepsy: a key link between structural and functional network disruptions. Brain. 2016; 139:2431–2440.

35. Bernhardt BC, Bernasconi N, Concha L, et al. Cortical thickness analysis in temporal lobe epilepsy: reproducibility and relation to outcome. Neurology. 2010; 74:1776–1784.

36. Larivière S, Weng Y, Vos de Wael R, et al. Functional connectome contractions in temporal lobe epilepsy: Microstructural underpinnings and predictors of surgical outcome. Epilepsia. 2020; 61:1221–1233.

37. Bernhardt BC, Kim H, Bernasconi N. Patterns of subregional mesiotemporal disease progression in temporal lobe epilepsy. Neurology. 2013; 81:1840–1847.

38. Bernhardt BC, Hong SJ, Bernasconi A, et al. Magnetic resonance imaging pattern learning in temporal lobe epilepsy: classification and prognostics. Ann Neurol. 2015; 77:436–446.

39. Zhang Z, Lu G, Zhong Y, et al. fMRI study of mesial temporal lobe epilepsy using amplitude of low-frequency fluctuation analysis. Hum Brain Mapp. 2010; 31:1851–1861.

40. Zeng H, Pizarro R, Nair VA, et al. Alterations in regional homogeneity of resting-state brain activity in mesial temporal lobe epilepsy. Epilepsia. 2013; 54:658–666.

41. Liao W, Zhang Z, Pan Z, et al. Default mode network abnormalities in mesial temporal lobe epilepsy: A study combining fMRI and DTI. Hum Brain Mapp. 2011; 32:883–895.

42. Remiszewski N, Bryant JE, Rutherford SE, et al. Contrasting case-control and normative reference approaches to capture clinically relevant structural brain abnormalities in patients with first-episode psychosis who are antipsychotic naive. JAMA Psychiatry. 2022; 79:1133–1138.

43. Bernhardt BC, Bernasconi A, Liu M, et al. The spectrum of structural and functional imaging abnormalities in temporal lobe epilepsy. Ann Neurol. 2016; 80:142–153.

44. Bernhardt BC, Fadaie F, Liu M, et al. Temporal lobe epilepsy: Hippocampal pathology modulates connectome topology and controllability. Neurology. 2019; 92:e2209–e2220.

45. Blanc F, Martinian L, Liagkouras I, et al. Investigation of widespread neocortical pathology associated with hippocampal sclerosis in epilepsy: A postmortem study. Epilepsia. 2011; 52:10-

46. Park HJ, Friston K. Structural and functional brain networks: from connections to cognition. Science. 2013; 342:1238411.

47. Lynn CW, Bassett DS. The physics of brain network structure, function and control. Nat Rev Phys. 2019; 1:318–332.

48. Suárez LE, Markello RD, Betzel RF, et al. Linking structure and function in macroscale brain networks. Trends Cogn Sci. 2020; 24:302–315.

49. Fotiadis P, Parkes L, Davis KA, et al. Structure–function coupling in macroscale human brain networks. Nat Rev Neurosci. 2024; 25:688–704.

50. Fornito A, Zalesky A, Breakspear M. The connectomics of brain disorders. Nat Rev Neurosci. 2015; 16:159–172.

51. Hansen JY, Shafiei G, Vogel JW, et al. Local molecular and global connectomic contributions to cross-disorder cortical abnormalities. Nat Commun. 2022; 13:4682.

52. Galovic M, van Dooren VQH, Postma T, et al. Progressive cortical thinning in patients with focal epilepsy. JAMA Neurol. 2019; 76:1230–1239.

53. Crossley NA, Mechelli A, Scott J, et al. The hubs of the human connectome are generally implicated in the anatomy of brain disorders. Brain. 2014; 137:2382–2395.

54. van den Heuvel MP, Sporns O. Network hubs in the human brain. Trends Cogn Sci. 2013; 17:683–696.

55. Fadaie F, Lee HM, Caldairou B, et al. Atypical functional connectome hierarchy impacts cognition in temporal lobe epilepsy. Epilepsia. 2021; 62:2589–2603.

56. Larivière S, Schaper FLWVJ, Royer J, et al. Brain networks for cortical atrophy and responsive neurostimulation in temporal lobe epilepsy. JAMA Neurol. 2024; 81:1199–1209.

57. Miron G, Müller PM, Hohmann L, et al. Cortical thickness patterns of cognitive impairment phenotypes in drug-resistant temporal lobe epilepsy. Ann Neurol. 2024; 95:984–997.

58. Stiso J, Khambhati AN, Menara T, et al. White matter network architecture guides direct electrical stimulation through optimal state transitions. Cell Rep. 2019; 28:2554-2566.e2557.

59. Tang E, Giusti C, Baum GL, et al. Developmental increases in white matter network controllability support a growing diversity of brain dynamics. Nat Commun. 2017; 8:1252.

60. Cataldi M, Avoli M, de Villers-Sidani E. Resting state networks in temporal lobe epilepsy. Epilepsia. 2013; 54:2048–2059.

61. Rodriguez-Cruces R, Velázquez-Pérez L, Rodríguez-Leyva I, et al. Association of white matter diffusion characteristics and cognitive deficits in temporal lobe epilepsy. Epilepsy Behav. 2018; 79:138–145.

62. Hatton SN, Huynh KH, Bonilha L, et al. White matter abnormalities across different epilepsy syndromes in adults: an ENIGMA-Epilepsy study. Brain. 2020; 143:2454–2473.

63. Gleichgerrcht E, Munsell BC, Alhusaini S, et al. Artificial intelligence for classification of temporal lobe epilepsy with ROI-level MRI data: A worldwide ENIGMA-Epilepsy study. Neuroimage Clin. 2021; 31:102765.

64. Erik K, Jun R, Allen JC, et al. Convolutional neural network algorithm to determine lateralization of seizure onset in patients with epilepsy. Neurology. 2023; 101:e324.

65. Abbasi B, Goldenholz DM. Machine learning applications in epilepsy. Epilepsia. 2019; 60:2037–2047.

66. Owen TW, Schroeder GM, Janiukstyte V, et al. MEG abnormalities and mechanisms of surgical failure in neocortical epilepsy. Epilepsia. 2023; 64:692–704.

67. Wilson W, Tehrani N, Pittman DJ, et al. Mapping interictal discharges using intracranial EEG-fMRI to predict postsurgical outcomes. Brain. 2024; 147:4157–4168.

68. Bonilha L, Helpern JA, Sainju R, et al. Presurgical connectome and postsurgical seizure control in temporal lobe epilepsy. Neurology. 2013; 81:1704.

69. Morgan VL, Rogers BP, Anderson AW, et al. Divergent network properties that predict early surgical failure versus late recurrence in temporal lobe epilepsy. J Neurosurg. 2020; 132:1324–1333.

70. Barba C, Rheims S, Minotti L, et al. Temporal plus epilepsy is a major determinant of temporal lobe surgery failures. Brain. 2016; 139:444–451.

71. Wu C, Sharan AD. Neurostimulation for the treatment of epilepsy: a review of acurrent surgical interventions. Neuromodulation. 2013; 16:10–24.

72. Jobst BC, Kapur R, Barkley GL, et al. Brain-responsive neurostimulation in patients with medically intractable seizures arising from eloquent and other neocortical areas. Epilepsia. 2017; 58:1005–1014.

73. Scheffer IE, Berkovic S, Capovilla G, et al. ILAE classification of the epilepsies: Position paper of the ILAE Commission for Classification and Terminology. Epilepsia. 2017; 58:512–521.

74. Jackson GD, Berkovic SF, Duncan JS, et al. Optimizing the diagnosis of hippocampal sclerosis using MR imaging. AJNR Am J Neuroradiol. 1993; 14:753.

75. Achten E, Boon P, De Poorter J, et al. An MR protocol for presurgical evaluation of patients with complex partial seizures of temporal lobe origin. AJNR Am J Neuroradiol. 1995; 16:1201.

76. Labate A, Ventura P, Gambardella A, et al. MRI evidence of mesial temporal sclerosis in sporadic “benign” temporal lobe epilepsy. Neurology. 2006; 66:562–565.

77. Rodriguez-Cruces R, Royer J, Herholz P, et al. Micapipe: a pipeline for multimodal neuroimaging and connectome analysis. NeuroImage. 2022; 263:119612.

78. Glasser MF, Coalson TS, Robinson EC, et al. A multi-modal parcellation of human cerebral cortex. Nature. 2016; 536:171–178.

79. Wehrheim MH, Faskowitz J, Schubert A-L, et al. Reliability of variability and complexity measures for task and task-free BOLD fMRI. Hum Brain Mapp. 2024; 45:e26778.

80. Baracchini G, Mišić B, Setton R, et al. Inter-regional BOLD signal variability is an organizational feature of functional brain networks. NeuroImage. 2021; 237:118149.

81. Schleifer CH, Chang SE, Amir CM, et al. Unique functional neuroimaging signatures of genetic versus clinical high risk for psychosis. Biol Psychiatry. 2024; 97:178–187.

82. Jiang L, Zuo X-N. Regional homogeneity: A multimodal, multiscale neuroimaging marker of the human connectome. Neuroscientist. 2015; 22:486–505.

83. Cole MW, Pathak S, Schneider W. Identifying the brain’s most globally connected regions. NeuroImage. 2010; 49:3132–3148.

84. Zuo XN, Ehmke R, Mennes M, et al. Network centrality in the human functional connectome. Cereb Cortex. 2012; 22:1862–1875.

85. Fortin JP, Cullen N, Sheline YI, et al. Harmonization of cortical thickness measurements across scanners and sites. NeuroImage. 2018; 167:104–120.

86. Johnson WE, Li C, Rabinovic A. Adjusting batch effects in microarray expression data using empirical Bayes methods. Biostatistics. 2007; 8:118–127.

87. Van Essen DC, Smith SM, Barch DM, et al. The WU-Minn Human Connectome Project: An overview. NeuroImage. 2013; 80:62–79.

88. Shafiei G, Bazinet V, Dadar M, et al. Network structure and transcriptomic vulnerability shape atrophy in frontotemporal dementia. Brain. 2022; 146:321–336.

89. Váša F, Mišić B. Null models in network neuroscience. Nat Rev Neurosci. 2022; 23:493–504.

90. Budescu DV. Dominance analysis: A new approach to the problem of relative importance of predictors in multiple regression. Psychol Bull. 1993; 114:542–551.

91. Chang CC, Lin CJ. LIBSVM: A library for support vector machines. ACM Trans Intell Syst Technol. 2011; 2:1–27.

