## Supplemental Methods and Results for "PERSONALIZED BIOMARKERS OF MULTISCALE FUNCTIONAL ALTERATIONS IN TEMPORAL LOBE EPILEPSY"

### MRI Acquisition

*MICA-MICs dataset*. Data were collected on a 3-T Siemens Magnetom Prisma-Fit scanner equipped with a 64-channel head coil, and included: (i) two T1-weighted scans (3D-MPRAGE, repetition time [TR] = 2300 ms, echo time [TE] = 3.14 ms, flip angle [FA] = 9°, field of view [FOV] = 256×256 $mm^2$, voxel size = 0.8×0.8×0.8 $mm^3$, matrix size = 320×320, 224 slices), (ii) a resting-state functional MRI (fMRI) scan (multiband accelerated 2D-BOLD echo-planar imaging (EPI), TR = 600 ms, TE = 30 ms, FA = 52°, FOV = 240×240 $mm^2$, voxel size = 3×3×3 $mm^3$, multi-band factor = 6, 48 slices, 700 volumes), and (iii) a multi-shell diffusion MRI scan (2D spin-echo EPI, TR = 3500 ms, TE = 64.40 ms, FA = 90°, FOV = 224×224 $mm^2$, voxel size = 1.6×1.6×1.6 $mm^3$, 3 b0 images, b-values = 300/700/2000 s/$mm^2$ with 10/40/90 diffusion directions). During the resting-state fMRI acquisition, participants were instructed to stay still, fixate a cross presented on the screen, and not to fall asleep.

*EpiC dataset*. Data were collected on a 3-T Philips Achieva scanner equipped with a 64-channel head coil, and included: (i) a T1-weighted scan (3D spoiled gradient-echo, TR = 8.1 ms, TE = 3.7 ms, FA = 8°, FOV = 256×256 $mm^2$, voxel size = 1×1×1 $mm^3$, 240 slices), (ii) a resting-state fMRI scan (gradient-echo EPI, TR = 2000 ms, TE = 30 ms, FA = 90°, voxel size = 2×2×3 $mm^3$, 34 slices, 200 volumes), and (iii) a diffusion MRI scan (2D EPI, TR = 11.86 s, TE = 64.3 ms, FOV = 256×256 $mm^2$, voxel size = 2×2×2 $mm^3$, 2 b0 images, b-value = 2000 s/$mm^2$, 60 diffusion directions). During the resting-state fMRI acquisition, participants were instructed to keep their eyes closed and not to fall asleep.

*Nanj dataset*. Data were collected on a 3-T Siemens Trio scanner equipped with a 32-channel head coil, and included: (i) a T1-weighted scan (3D-MPRAGE, TR = 2300 ms, TE = 2.98 ms, FA = 9°, FOV = 256×256 $mm^2$, voxel size = 0.5×0.5×1 $mm^3$), (ii) a resting-state fMRI scan (2D gradient-echo EPI, TR = 2000 ms, TE = 30 ms, FA = 90°, FOV = 240×240 $mm^2$, voxel size = 3.75×3.75×4 $mm^3$, 30 slices, 255 volumes), and (iii) a diffusion MRI scan (2D spin-echo EPI, TR = 6100 ms, TE = 93 ms, FA = 90°, FOV = 240×240 $mm^2$, voxel size = 0.94×0.94×3 $mm^3$, 4 b0 images, b-value = 1000 s/$mm^2$, 120 diffusion directions). During the resting-state fMRI acquisition, participants were instructed to keep their eyes closed and not to fall asleep.

*NOEL dataset*. Data were collected on a 3-T Siemens Trio scanner equipped with a 32-channel head coil, and included: (i) a T1-weighted scan (3D-MPRAGE, TR = 2300 ms, TE = 2.98 ms, FA = 9°, voxel size = 1×1×1 $mm^3$), (ii) a resting-state fMRI scan (2D gradient-echo EPI, TR = 2020 ms, TE = 30 ms, FA = 90°, voxel size = 4×4×4 $mm^3$, 34 slices, 150 volumes), and (iii) a diffusion MRI scan (2D twice-refocused EPI, TR = 8400 ms, TE = 90 ms, FA = 90°, voxel size = 2×2×3 $mm^3$, 63 slices, 1 b0 images, b-value = 1000 s/$mm^2$, 64 diffusion directions). During the resting-state fMRI acquisition, participants were instructed to keep their eyes closed and not to fall asleep.

### MRI Processing

Multimodal MRI data were preprocessed using *micapipe* (v0.2.3; https://micapipe.readthedocs.io/),[1] an openly accessible multimodal MRI pipeline that integrates AFNI, FSL, FreeSurfer, ANTs, MRtrix, and Workbench.[2-6] T1-weighted MRI data were de-obliqued, reoriented to standard orientation (LPI: left to right, posterior to anterior, and inferior to superior), linearly co-registered, corrected for intensity non-uniformity, intensity normalized, skull stripped, and submitted to FreeSurfer 6.0 to extract models of the inner and outer cortical interfaces. Segmentation errors were manually corrected. Subject-specific cortical thickness was measured as Euclidean distance between corresponding pial and white matter vertices, and registered to the Conte69 template surface (~32k vertices/hemisphere).[7]

Diffusion MRI data were denoised, and corrected for susceptibility distortions, head motion, and eddy currents using MRtrix3 (http://www.mrtrix.org/).[6] A Laplacian potential field was used to guide the placement of a superior white matter (SWM) surface, targeting a depth of ~2 mm beneath the graywhite matter boundary.[8,9] Diffusion features, fractional anisotropy (FA) and mean diffusivity (MD), which serve as surrogates of fiber architecture and tissue microstructure, were linearly interpolated along the SWM surface and registered to the Conte69 template surface.

Resting-state fMRI preprocessing included discarding the first five volumes, reorientation, slice-timing correction, and head motion correction. Nuisance signals were removed using an in-house trained ICA-FIX classifier.[10] Preprocessed time series were non-linearly registered to native FreeSurfer space using boundary-based registration and mapped to the Conte69 template surface.

Surface-based maps, including cortical thickness, FA, MD, and resting-state fMRI time series, were spatially smoothed using a Gaussian kernel with a full-width-at-half-maximum (FWHM) of 10 mm. Finally, vertex-wise cortical maps were parcellated using the HCPMMP1.0 atlas (or Glasser atlas),[11] a multimodal parcellation of the cerebral cortex with 180 homologous parcels per hemisphere.

### REFERENCES


1. Rodriguez-Cruces R, Royer J, Herholz P, et al. Micapipe: a pipeline for multimodal neuroimaging and connectome analysis. *NeuroImage*. 2022; 263:119612.
2. Cox RW. AFNI: Software for analysis and visualization of functional magnetic resonance neuroimages. *Comput Biomed Res*. 1996; 29:162-173.
3. Jenkinson M, Beckmann CF, Behrens TEJ, et al. FSL. *NeuroImage*. 2012; 62:782-790.
4. Fischl B. FreeSurfer. *NeuroImage*. 2012; 62:774-781.
5. Avants BB, Epstein CL, Grossman M, et al. Symmetric diffeomorphic image registration with cross-correlation: Evaluating automated labeling of elderly and neurodegenerative brain. *Med Image Anal*. 2008; 12:26-41.
6. Tournier JD, Smith R, Raffelt D, et al. MRtrix3: A fast, flexible and open software framework for medical image processing and visualisation. *NeuroImage*. 2019; 202:116137.
7. Van Essen DC, Glasser MF, Dierker DL, et al. Parcellations and hemispheric asymmetries of human cerebral cortex analyzed on surface-based atlases. *Cereb Cortex*. 2012; 22:2241-2262.
8. Liu M, Bernhardt BC, Hong SJ, et al. The superficial white matter in temporal lobe epilepsy: a key link between structural and functional network disruptions. *Brain*. 2016; 139:2431-2440.
9. Larivière S, Weng Y, Vos de Wael R, et al. Functional connectome contractions in temporal lobe epilepsy: Microstructural underpinnings and predictors of surgical outcome. *Epilepsia*. 2020; 61:1221-1233.
10. Salimi-Khorshidi G, Douaud G, Beckmann CF, et al. Automatic denoising of functional MRI data: Combining independent component analysis and hierarchical fusion of classifiers. *NeuroImage*. 2014; 90:449-468.
11. Glasser MF, Coalson TS, Robinson EC, et al. A multi-modal parcellation of human cerebral cortex. *Nature*. 2016; 536:171-178.

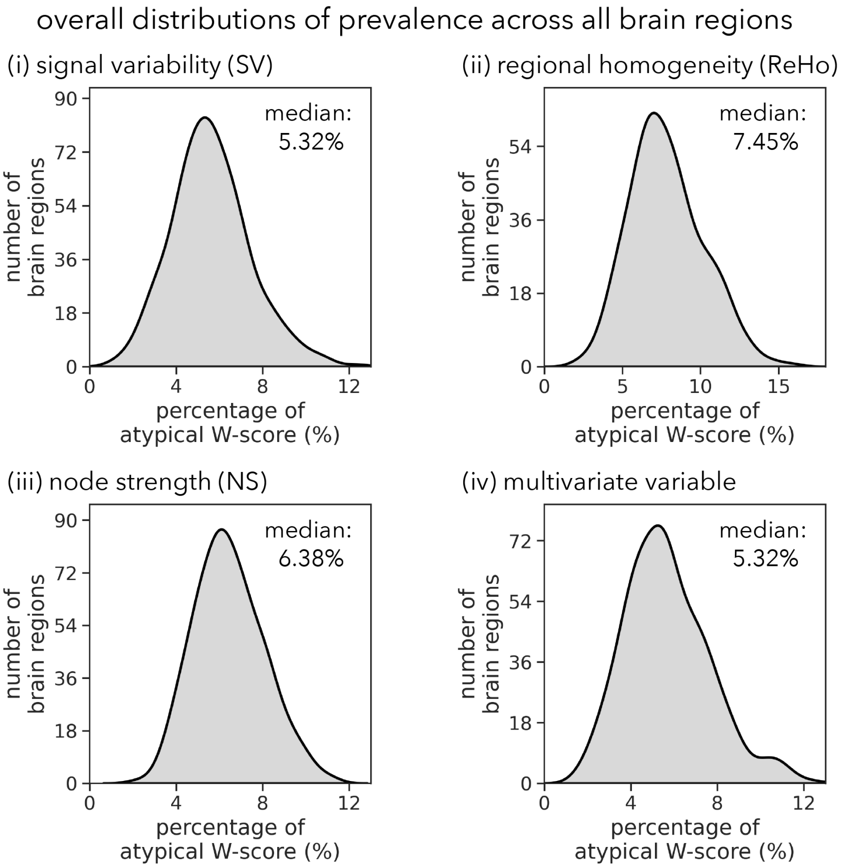


**Figure S1**: The overall distributions of the proportion of extreme deviations in TLE patients across all brain regions.

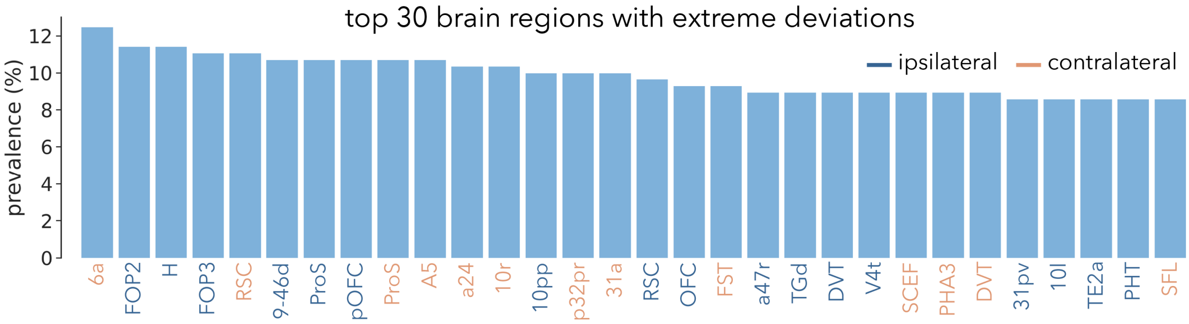


**Figure S2**: The 30 brain regions with the highest proportion of extreme deviations in TLE patients.

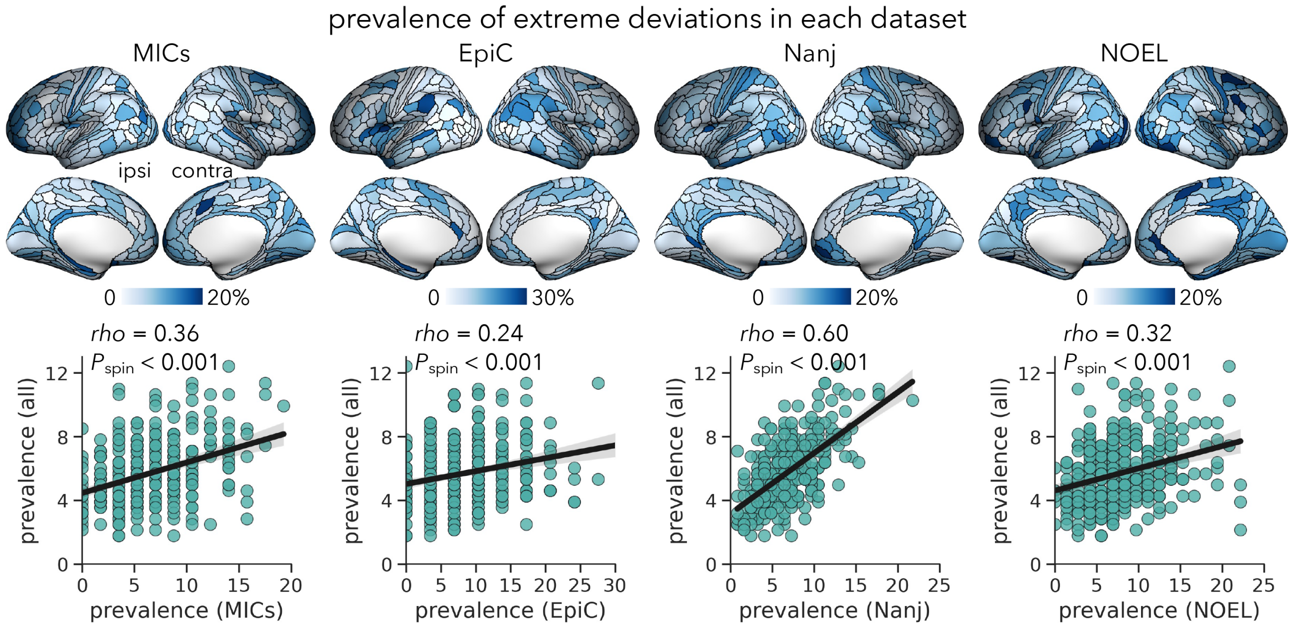


**Figure S3**: **Region-specific prevalence of extreme deviations of TLE patients in each dataset**. *Top*: Proportion of TLE patients with *W*-scores exceeding ±1.96 in each brain region. *Bottom*: Spatial correlations between each dataset's deviation prevalence map (x-axis) and the four-dataset pooled deviation prevalence map (y-axis, **Figure 2B**), with each dot representing a single brain region. Statistical significance (*i.e.*, $P_{spin}$) of correlation coefficients are determined using spin permutation tests with 5,000 iterations. ipsi = ipsilateral; contra = contralateral.

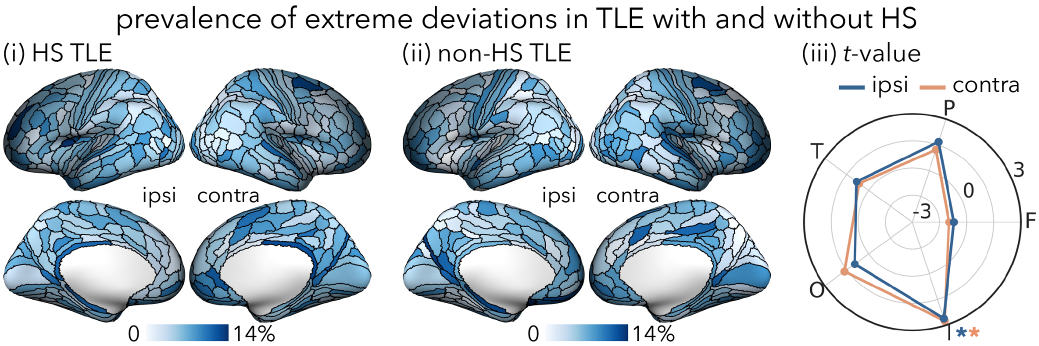

**Figure S4**: **Region-specific prevalence of extreme deviations in TLE patients with and without hippocampal sclerosis (HS)**. Proportion of patients with extreme deviations in each brain region in (*i*) HS and (*ii*) non-HS TLE subgroups separately. (iii) Differences in the number of extreme deviations between HS and non-HS patients in each lobe. * $P_{FDR} < 0.05$. F = frontal; P = parietal; T = temporal; O = occipital; I = insula; ipsi = ipsilateral; contra = contralateral.

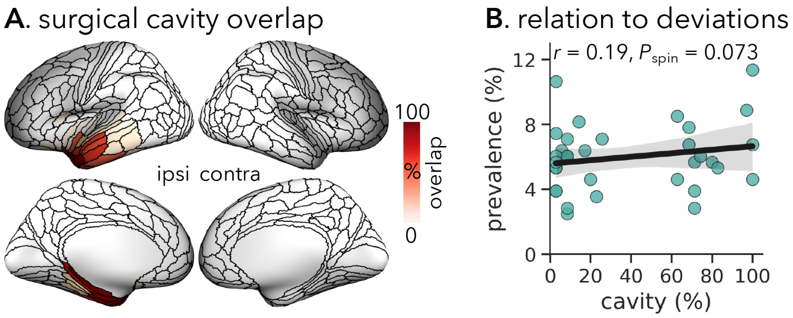


**Figure S5**: **Spatial correlation between the overlap of surgical cavities and TLE-related functional deviations**. **(A)** The overlap map of surgical cavities segmented from pre- and post-surgical T1-weighted MRIs across 35 TLE patients. **(B)** There is a positive association between the overlap map of surgical cavities and the proportion of extreme deviations within the resected brain regions after spin permutation tests with 5,000 iterations. ipsi = ipsilateral; contra = contralateral.

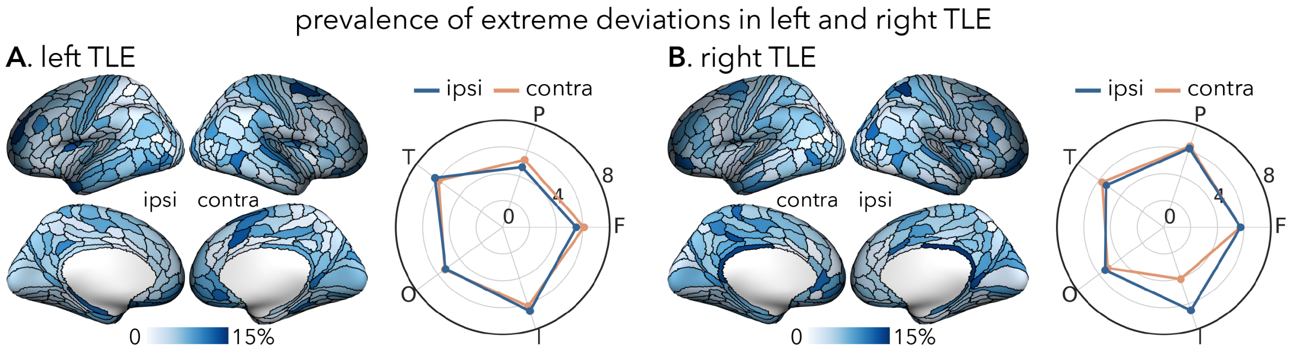


**Figure S6**: **Region-specific prevalence of extreme deviations in (A) left and (B) right TLE subgroups**. Proportion of patients with extreme deviations in each brain region and lobe in left and right TLE subgroups separately. F = frontal; P = parietal; T = temporal; O = occipital; I = insula; ipsi = ipsilateral; contra = contralateral.

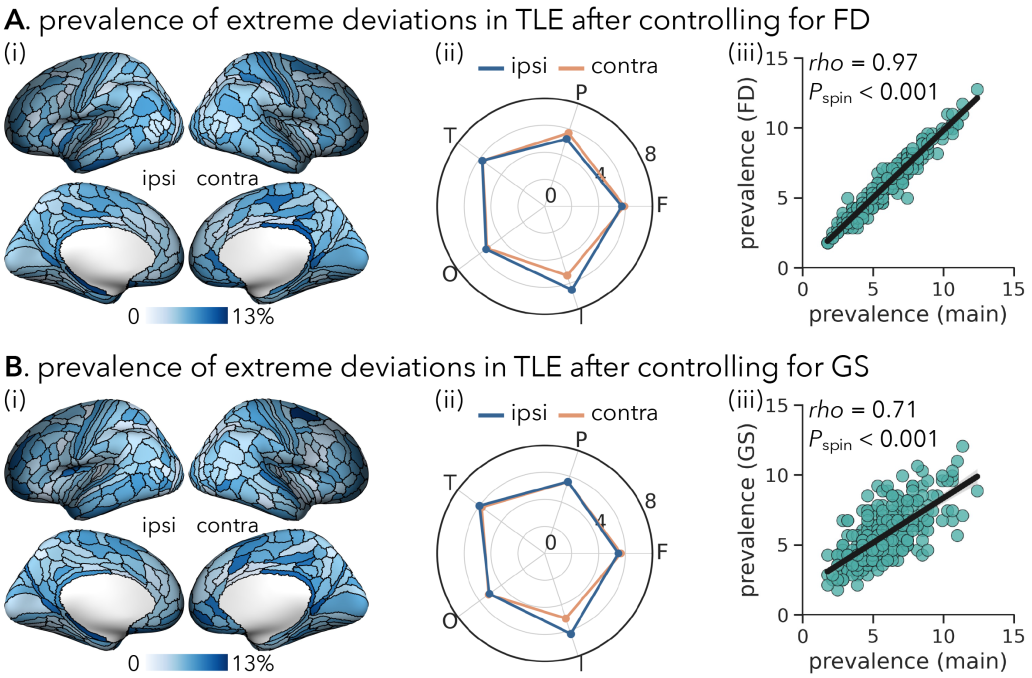


**Figure S7**: **Region-specific prevalence of extreme deviations in TLE patients after controlling for (A) FD or (B) GS**. (*i*) Proportion of patients with extreme deviations in each brain region when additionally controlling for head motion (calculating as framewise displacement (FD)) in rs-fMRI scans or global mean signal (GS). (*ii*) Mean proportion of extreme deviations in each lobe. (*iii*) Spatial correlations between the deviation prevalence patterns before (x-axis, **Figure 2B**) and after (y-axis) controlling for FD or GS. F = frontal; P = parietal; T = temporal; O = occipital; I = insula; ipsi = ipsilateral; contra = contralateral.
